## Supplementary information for "Labeling of Nascent RNA in *C. elegans* Intestine"

### Table of contents

- Supplementary File 1 (detailed protocol)
- Supplementary Table 1

### Labeling of Nascent RNA in *C. elegans* Intestine Cells

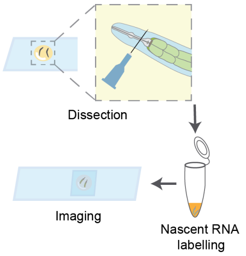

RESERVED DOI:
10.17504/protocols.io.rm7vzqr38vx1/v1

Omid Gholamalamdari[1], Stephanie C. Weber[1,2]

[1]Department of Biology, McGill University; [2]Department of Physics, McGill University

Omid Gholamalamdari
McGill University

**Protocol Info:** Omid Gholamalamdari, Stephanie C. Weber . Labeling of Nascent RNA in C. elegans Intestine Cells. **protocols.io** **https://protocols.io/view/labeling-of-nascent-rna-in-c-elegans-intestine-cel-gyxibxxkf**

**Created:** May 06, 2025

**Last Modified:** September 08, 2025

**Protocol Integer ID:** 217802

**Keywords:** RNA labelling, nascent RNA, C. elegans, intestine, click chemistry, microscopy,

**Funders Acknowledgements:**
**Canadian Institutes of Health Research**
**Grant ID: 159580**

#### Abstract

We present a novel protocol for metabolic labeling of nascent RNA in the *C. elegans* intestine, overcoming the challenge of RNA analogs' inability to penetrate the worm cuticle. This method involves dissecting worms to extrude their intestines and performing nascent RNA labeling in a tube. While optimized for imaging applications, the protocol can be adapted for other molecular techniques, such as sequencing or RT-qPCR, enabling quantitative analysis of nascent transcripts. This approach offers a powerful tool for investigating transcriptional regulation and RNA metabolism in the context of *C. elegans* physiology and disease models.

#### Guidelines

Stage your worms according to the needs of your experiment.
This protocol can be performed on L4 larvae, young adults, and aged adults.

#### Materials

1. Positively charged microscope slides, for dissection (Fisherbrand 22-037-246)
2. Meiosis Media (MM), adapted from Laband et al. 2018

| Reagent | Amount |
|---|---|
| Libovitz's L-15 media | 6 mL |
| 1 M HEPES, pH 7.5 | 125 µL |
| Heat-inactivated FBS | 2 mL |
| Inulin | 5 mg |
| Nuclease-free ddH2O | to 10 mL |
| Filter sterilize. Aliquot and freeze. | |

3. Dissection Media: 196 µL MM + 4 µL 2% (w/v) Tetramisole
4. Syringe needle, 25G
5. Aspirator tube assemblies (Sigma cat A5177)
6. Microcapillary tube (50 µL calibrated pipet; Drummond Scientific Company cat 2-000-050)
   - Flame, pull, and break the capillary tubes to make two mouth pipets
   - Briefly flame the broken tip, so that the edges become smooth
   - One with wider opening, for handling intestines
   - One with very narrow opening, for handling liquids
7. Siliconized centrifuge tubes (BIO PLAS cat 4165SL)
8. RNA Labeling Media: 18 µL MM + 2 µL 10 mM EU
9. M9 buffer: 3 g $KH_2PO_4$, 6 g $Na_2HPO_4$, 5 g NaCl, 1 ml 1 M $MgSO_4$, $H_2O$ to 1 L. Sterilize by autoclaving.
10. 60% isopropanol
11. Washing Solution: 1X PBS + 0.1 % Triton X-100 (referred to **PBS-Tx** in the protocol)
12. Permeabilizing Solution: 1X PBS + 0.5% Triton X-100
13. Click chemistry components
   - Fluorophore-azide of choice (1 mM); aliquot and freeze
   - $CuSO_4$ (20 mM); keep at room temperature
   - THPTA (100 mM); aliquot and freeze
   - Vitamin C (60 mg/ml); **prepare fresh**
14. Mounting Media
   - Mix reagents 1-3 in 15 mL conical tube and heat at 65˚C for 10 minutes, or until n-propyl gallate dissolves.
   - Add glycerol and DAPI. Aliquot and store at -20˚C.

| Reagent | Amount |
|---|---|
| n-propyl gallate | 800 mg |
| 1 M Tris, pH 9 | 0.3 mL |
| Nuclease-free ddH2O | 2.7 mL |
| Glycerol | 7 mL |
| DAPI (2.5 mg/mL) | 8 µL |

15. Microscope slides, for mounting

16. Cover slips, #1.5 22×22 mm

#### Before start

On the day of the experiment, prepare the Dissection Media and RNA Labeling Media. Keep at room temperature.

#### Dissection of intestine tissue

15m

1 On a positively charged slide, place 4 drops of 10 µL Dissection Media (DM).

- For better extrusion of intestine from older worms (>= Day 5 adults), use 0.02% Tetramisole in DM.

2 Using a worm pick, transfer 5 worms from an NGM plate to each of the DM drops.

- Set a timer for 10 minutes.

10m

3 Working quickly on a dissecting scope, use a syringe needle to cut each worm between the pharynx and the mouth.

- We recommend holding a syringe needle between your thumb and index finger and stabilize it with a finger from the opposite hand.

Once the intestine extrudes, detach it from the rest of the worm by cutting through the mid-line of the worm's anterior-posterior axis (i.e. in the vicinity of the vuvla).

- Change the needle frequently (~every 20 dissection).
- Needles have minor differences in their shape and sharpness. If cutting is challenging, simply changing the needle might help.

4 When the dissection is over or the timer is up, transfer the intestines using the mouth pipet with large opening to a siliconized tube.

- Due to the large opening of the mouth pipet and small volume of the sample, it is necessary to use the mouth pipet in a controlled way.
- To avoid sudden suction and sample loss, consciously switch breathing between nose and mouth. (i.e. through your mouth while not suctioning and through your nose while suctioning)
- Adding more DM ( 10 µL ) to the intestines before mouth pipetting can be helpful.
- Marker-written labels can easily wipe off the siliconized tube. To prevent this, label tube on the frosted side or with a sticker on the lid.

5 Spin down the intestines, 1500 x g, Room temperature, 00:00:30 .

**CRITICAL STEP:** Higher g-forces damage the cells and may affect transcription.

#### Metabolic labeling of nascent RNA

20m

6 Using the dissecting scope and mouth pipet with narrow opening, remove the Dissection Media.

7 Add 50 µL of RNA Labeling Media with a P200 pipet and gently pipet up and down twice.

8 Incubate for 00:05:00 with shaking at 1300 rpm, Room temperature .

5m

9 Add 500 µL of M9 buffer to dilute the EU in the RNA Labeling Media and stop the labeling.

10 Spin down the intestines at 1500 x g, Room temperature, 00:01:00

11 Under the dissecting scope, remove the M9 buffer using a P1000 pipet and leave around 50 µL behind.

- Centrifuge for an additional minute if the intestines have not pelleted completely.

12 Fix the samples by adding 500 µL of room temperature 60% isopropanol. Incubate for 00:10:00 .

13 Samples can be kept at -20 °C for up to two weeks.

#### Click chemistry

2h

14 Wash the samples with PBS-Tx.

- From this point, a wash is defined as adding 500 µL of a solution, centrifuging at 8000 x g, Room temperature, 00:01:00 , and removing the wash solution with a P1000 pipet under the dissecting scope.

15 **CRITICAL STEP:**

Permeabilize the intestines with 500 µL 1X PBS + 0.5% Triton X-100 for 00:05:00 .

- Longer exposure of intestines to Permeabilizing Solution results in greater extraction of cellular material. Therefore, sample handling time for removing the Permeabilizing Solution may result in extraction of cellular RNA.
- Hence, performing this step with a large number of samples is not recommended.

16 Wash twice with PBS-Tx.

17 Prepare the Vitamin C solution fresh: Measure 30 to 60 mg of Vitamin C in a 1.5 mL tube and accordingly add nuclease-free dd$H_2O$ to make [M] 60 mg/mL Vitamin C solution.

- It is **critical** to make Vitamin C solution fresh

18 Prepare the Click Chemistry Solution as below, up to 15 minutes before use.

18.1

| Reagent | Stock concentration | For 1 sample | For 6 samples |
|---|---|---|---|
| PBS-Tx | 1X | 83 µL | 498 |
| Fluorophore-azide of choice | 1 mM | 2 µL | 12 |
| CuSO4 | 20 mM | 5 µL | 30 |
| THPTA | 100 mM | 5 µL | 30 |
| Vitamin C | 60 mg/ml | 5 µL | 30 |
| Total | | 100 µL | 600 |

**Make Vitamin C fresh!**

19 Incubate 1300 rpm, Room temperature , 00:30:00 in the dark 30m

20 Wash three times with PBS-Tx.

#### Mounting 30m

21 Spin down samples at 8000 x g, Room temperature, 00:01:00 .

22 Carefully, remove all but 100 µL of PBS-Tx using a P1000 pipet.

23 Carefully, remove all but 25 µL of PBS-Tx using a P200 pipet.

24 Remove the rest of the PBS-Tx using the liquid handling mouth pipet.

25 Add 50 µL of Mounting Media and gently pipet several times with P100.

- If not mixed well, the intestines will float on top of Mounting Media after centrifugation.

26 Spin down samples at 8000 x g, 00:01:00 .

27 On a normal glass slide, use a pap-pen to draw a square of ~1.5 cm side length.

Centered inside the square, draw a circle of ~1 cm diameter.

- This pattern is necessary to prevent intestines from moving to the edges when the cover slip is applied.
- Intestines will be loaded inside the circle, and extra Mounting Media will be added to the corners between the circle and square.
- Once the coverslip is placed, the outer area will create a barrier. Once all the mounting media is fused, this prevents the intestines from moving to the edges of the coverslip.
- This drawing should be done with swift motions. Excessive amounts of pap-pen marks may hinder the fusing of mounting media from outer and inner areas.

28 Looking under dissecting scope, pipet 15 µL of the sample and mount in the center of the circle on the slide.

- Try to get all of the intestines.

29 Take another 15 µL of Mounting Media from the same tube (try to take remaining intestines if anything is left) and place 4 drops of equal volume in the corners.

The slide should look like this in the end:

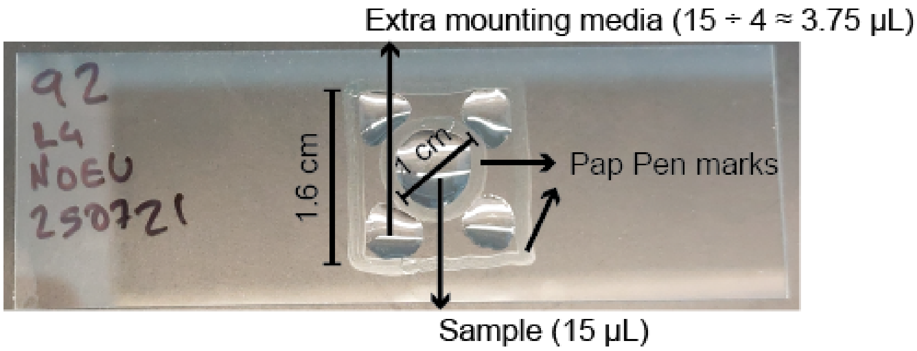

30 Carefully drop a cover slip on top. It is important that the cover slip touches the central circle (sample) first.

- Bubbles may form, but they do not interfere with subsequent imaging.

31 Seal with nail polish.

32 Let the nail polish cure at room temperature overnight in the dark.

33 Store the slides at 4°C and image within one week.

#### Protocol references

Laband, Kimberley, Benjamin Lacroix, Frances Edwards, Julie C. Canman, and Julien Dumont. "Live Imaging of *C. elegans* Oocytes and Early Embryos." In *Methods in Cell Biology*, edited by Helder Maiato and Melina Schuh, 145:217–36. Mitosis and Meiosis Part B. Academic Press, 2018. https://doi.org/10.1016/bs.mcb.2018.03.025.

#### Acknowledgements

We thank Laeya Baldini, Eric Cheng, Abigail Gerhold, and Réda Zellag for technical suggestions that led to the optimization of this protocol.

### Supplementary Table 1

| Name | Description | Genotype | Additional Information | Source |
|---|---|---|---|---|
| WLW92 | nucl-1::split-gfp | muIs253 [eft-3p::sfGFP1-10::unc-54 3'UTR + Cbr-unc-119(+)] II; nucl-1(sam132[NUCL1::M3]) IV | cross between DUP243 and CF4587 | This study |
| WLW93 | garr-1::split-gfp | | cross between DUP250 and CF4587 | This study |
| WLW3 | rpoa-2::gfp | ptnsIs053[pCPB155;pCPB157-No1] | | |
| WLW2 | dao-5::gfp | ptnIs050[dao-5::gfp] | | |
